# Accelerating Wound Healing Through Deep Reinforcement Learning: A Data-Driven Approach to Optimal Treatment

**DOI:** 10.1101/2024.12.18.629218

**Authors:** Fan Lu, Ksenia Zlobina, Prahbat Baniya, Houpu Li, Nicholas Rondoni, Narges Asefifeyzabadi, Wan Shen Hee, Maryam Tebyani, Kaelan Schorger, Celeste Franco, Michelle Bagood, Mircea Teodorescu, Marco Rolandi, Rivkah Isseroff, Marcella Gomez

**Affiliations:** University of California, Santa Cruz, 1156 High St, Santa Cruz, CA 95064, USA; University of California, Davis, 1 Shields Ave, Davis, CA 95616, USA; Clemson University, South Carolina 29634 USA

**Keywords:** Wound healing, Deep Learning, Deep Reinforcement Learning

## Abstract

Advancements in bioelectronic sensors and actuators have paved the way for real-time monitoring and control of wound healing progression. Real-time monitoring allows for precise adjustment in treatment strategies that align with an individual’s unique biological response. However, due to the complexities of human-drug interactions and a lack of predictive models it is challenging to determine just how one should adjust drug dosage to achieve the desired biological response. This work proposes an adaptive closed-loop control framework that integrates deep learning, optimal control, and reinforcement learning to update treatment strategies in real-time with the goal of accelerating wound closure. The proposed approach eliminates the need for mathematical modeling of complex nonlinear wound healing dynamics. We demonstrate the convergence of the controller via an *in silico* experimental setup, where the proposed approach successfully accelerates the wound healing process by 17.71%. Finally, we share the experimental setup and results of an *in vivo* implementation to highlight the translational potential of our work. Our data-driven model estimates a 40% acceleration in wound closure.

## Introduction

Wound healing is a dynamic and continuous process that involves nonlinear interactions across different cell types (platelets, neutrophils, macrophages, myofibroblasts, fibroblasts, keratinocytes, and others) and bio-molecules (blood coagulation factors, pro- and anti-inflammatory cytokines, polymers and enzymes of extracellular matrix) (1). These nonlinear processes can be modulated by different treatments administrated on the wound, influencing the healing process. In this context, personalized precision treatments have emerged as a vital research area in modern health care, driven by recent advancements in artificial intelligence (2–4). The need for precision treatment arises from the fact that patients often exhibit varied responses to the same medication, a phenomenon that arises from both molecular differences between individuals and variations within the same patient over time (5). Personalized treatment aims to identify the most effective drug types, dosages, and timing of administration tailored to each patient’s unique responses, leveraging experimental data and statistical analysis (6).

In this work, we explore two treatment modalities in this paper: electric field (EF) therapy and fluoxetine (Flx), both of which have demonstrated efficacy in accelerating wound healing. Endogenous electric fields facilitate healing by promoting the migration of epidermal stem cells, which are vital for tissue repair (7). Previous studies have shown that fluoxetine, when administered either systemically or topically, enhances wound healing in both diabetic and non-diabetic rodent models (8). Moreover, fluoxetine has been found to inhibit pathogen growth, reduce biofilm formation, and limit the spread of infection in rodents (9).

Determining the optimal intensity of the electric field, fluoxetine dosage, and the timing for switching between different types of treatments presents a significant challenge, particularly when accounting for the nonlinear dynamics of cell migration induced by electric fields, drug metabolism, and the biological interventions targeted by the medication. While mathematical models can help elucidate the mechanisms of electric field and drug distribution (10, 11), the complexity of biological systems and individual variability can undermine the reliability of model-based controllers.

Even with accurate models, the inherent nonlinearity complicates the task of ensuring the optimality and safety of prescribed treatment strategies for wound care. Conversely, the study of linear systems is well established, providing scalable methods for designing, analyzing, controlling, and optimizing such systems (12, 13).

However, establishing a reliable mapping from nonlinear systems to linear models remains challenging. The Koopman operator theory, as discussed in foundational works (14–16), offers a promising approach by allowing a nonlinear system to be represented as an infinite-dimensional linear system. Yet, the optimization of this framework typically operates within functional space, which can be impractical in realworld applications and does not account for control inputs in nonlinear systems.

Recent advances have sought to generalize Koopman operator theory for controlling nonlinear systems (17–20). These extensions provide finite-dimensional function approximations that can facilitate the tractable control of nonlinear systems. This motivated us to design a deep neural network-based algorithm to learn a mapping from an unknown non-linear system to its linear representation, akin to the objectives of the Koopman operator. Our work builds upon previous research (19–21). The major difference is that our approach addresses overfitting during training and does not rely on prior knowledge of a control-affine matrix. In contrast, we mitigate these challenges through the integration of optimal control and deep reinforcement learning (DRL).

The main contributions of this paper include: (i) We propose an adaptive closed-loop control framework using deep learning, optimal control, and reinforcement learning to enhance wound healing, as schematized in Fig. 1. This framework eliminates the need for mathematical modeling of non-linear dynamics or the mechanism by which a treatment effects wound healing processes. (ii) We extend the DeepMapper method from (21), eliminating the need for a control-affine matrix. Our results demonstrate its ability to learn a linear representation of nonlinear wound healing dynamics from a time-series of images monitoring wound closure *in-vivo*. By leveraging optimal control and the decoder mechanism within DeepMapper, we derive an optimal reference signal that the DRL agent uses to generate real-time treatment strategies. This approach not only improves the accuracy of the linear representation in modeling wound dynamics under optimal treatments but also enhances the efficiency of the reinforcement learning agent. (iii) We validate the efficacy of our approach with experimental data. Simulations of nonlinear healing dynamics with adaptive interventions show that the proposed method accelerated wound healing by 17.71% compared to normal healing process. Additionally, the proposed framework has successfully been implemented in *in-vivo* porcine experiments, demonstrating our method’s potential for translation to clinical settings (22). The proposed framework showcases the significant potential for expediting wound healing by effectively integrating optimal control and data-driven methods. By leveraging advanced algorithms that adapt in real-time to changing conditions, this system offers a more accurate and reliable means of promoting faster recovery without relying on the limitations of conventional models.

**Fig. 1.**
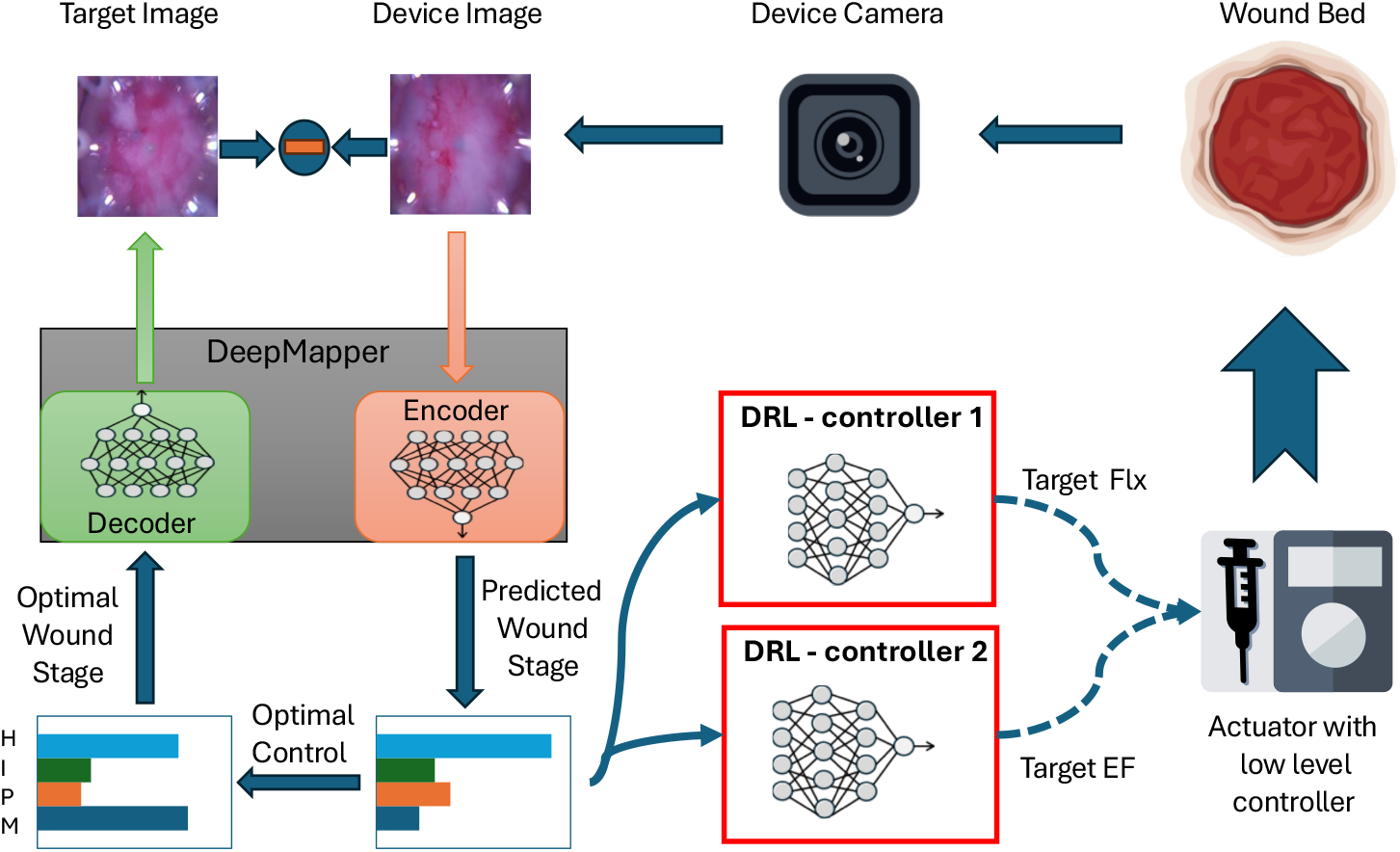
The DRL-based closed-loop control pipeline for accelerating wound healing involves three steps. First, a wound image captured by a camera is fed into the DeepMapper, which predicts the wound stage. Second, the deep reinforcement learning (DRL) agents generate treatment strategies. Third, electric field or fluoxetine is delivered to the wound through an actuator.

## Approach

The control and optimization of wound-healing dynamics, particularly when devising the most effective treatment strategy, is challenging due to their inherently nonlinear nature. Previous work (19–21) has focused on finding linear representations of nonlinear systems, from which optimal control can be derived. However, these approaches often require detailed knowledge of the control effects on the learned linear systems, such as a control-affine matrix, as in (21). In practice, obtaining this information is often difficult, making it challenging to translate the learned linear system into actionable treatment parameters, such as drug dosage or electric field strength.

To address this gap, we employ a leader-follower strategy, commonly used in robotic control [2], by leveraging a decoder alongside a deep reinforcement learning (DRL) controller. This is made possible by DeepMapper, which learns both a linear representation of the nonlinear system and uses it to compute an optimal control strategy, bridging the gap between theoretical models and practical treatment parameters.

### DeepMapper: linearization of nonlinear wound-healing dynamics

Consider a nonlinear state space model in discrete time,

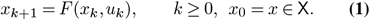

The state space X is a closed subset of ℝ^*n*^, and the input (or action) space ⋃ is finite, with cardinality *n*_⋃_ := | ⋃|, and where *F* : X *×* ⋃ *→* X. We may have state-dependent constraints, so that for each *x* ∈ X there is a set ⋃(*x*) ⊂ ⋃ for which *u*_*k*_ is constrained to *U* (*x*_*k*_) for each k.

Assume that there exists one unique equilibrium *x*^*e*^, which represent the full closure of the wound. This paper concerns finding an optimal control strategy *u*^⋆^ ∈ ϕ^⋆^(*x*) such that the time taken from any initial state *x* to an equilibrium *x*^*e*^ is minimized.

As discussed in (11), the function *F* can be an unmanageable nonlinear function that defines different cell transitions during wound healing. The nonlinear state *x* associated with wound healing surveyed in the literature may include variables such as pH, temperature (23–28), or visual representations captured through images of the wound (29). We assume that *x* can be measured by some sensor, but *F* is unknown to the control algorithm.

Solving for the optimal control policy *u*^⋆^ ∈ ϕ^⋆^(*x*) is often difficult, particularly when the dynamics evolve nonlinearly. Given that extensive research and literature has been dedicated to the study of linear systems, encompassing scalable design, analysis, control, and optimization (12, 30), we propose the utilization of a deep learning approach to model a linear system that best approximates the behavior of the underlying nonlinear system to guide control efforts. This builds on our previous work (21) for a more generalized approach. The major differences are the following: 1. in this work, we do not assume knowledge of a control affine matrix. we formulate the problem in discrete time, thus removing the need for calculating the Jacobian matrix of a deep neural network, and improving the efficiency of the algorithm.

Note that Eq. (1) defines a control-affine system. As discussed in (11, 19), the decoupling of the states and inputs allows us to find a transformation of the states alone:

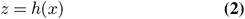

where *x* ∈ ℝ^*n*^is the state that evolves subject to nonlinear dynamics, *h* : ℝ^*n*^ *→* ℝ^*d*^ is a function that maps the nonlinear state *x* to linear state *z*, and *z* ∈ ℝ^*d*^ evolves linearly:

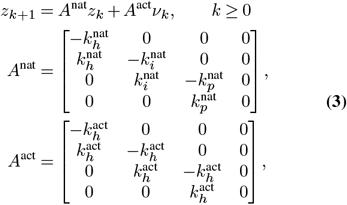

where *z*_0_ := *z* = [*H, I, P, M*]^⊺^ ∈ [0, 1]^4^, *A*^nat^, *A*^act^ ∈ [0, 1]^4*×*4^ contains values that control the velocities of transitions from homeostasis *H* to inflammation *I*, from inflammation to proliferation *P*, and from proliferation to maturation *M*.

Assume that *A*^nat^ + *A*^act^ ∈ [0, 1]^4^, with initial condition *z* = [1, 0, 0, 0]^⊺^, there exists one unique equilibrium in the linear dynamics: *z*^*e*^ = [0, 0, 0, 1]^⊺^, which represent full closure of the wound (100% of maturation). Without loss of generality, we assume that *z*^*e*^ = *h*(*x*^*e*^). Thus, minimizing the time taken from any initial state *x* to an equilibrium *x*^*e*^ is equivalent to minimizing:

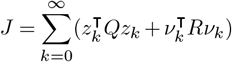

with 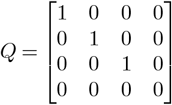 and 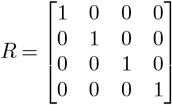.

The optimal control problem for the linear dynamcis Eq. (3) can be solved via the discrete time algebraic Riccati equation, giving the optimal control law:

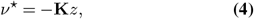

referred to as the Linear Quadratic Regulator (LQR) (30). The optimal control law in Eq. (4) only informs the required transition rates between wound healing stages to minimize the time for maturation to reach 100%. However, it does not give any information on what treatments should be applied to the wound, such as dosage of drug. As such, we need a mechanism that can map the linear state *z* to the nonlinear state *x* to inform real treatment *u*. In this paper, we adopt the updated DeepMapper architecture proposed in (31) where we employ three types of deep neural networks to parameterize the transformation function *h*, an inverse transformation function *h*^−1^ : ℝ^*d*^ *→* ℝ^*n*^, and the two matrices *A*^nat^ and *A*^act^ in Eq. (3).

Similar to (31), there are three high-level requirements for the neural networks, corresponding to three types of loss function used in training:

### Intrinsic coordinates that are useful for reconstruction

We seek to identify a few intrinsic coordinates *z* = *h*(*x*) where the dynamics evolve, along with the inverse *x* = *h*^−1^(*z*) so that the state *x* may be recovered. Reconstruction accuracy of the auto-encoder is achieved via minimizing the loss:

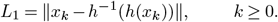

with ∥ · ∥ the mean-squared error, averaging over dimension then number of examples.

### Future state prediction

The intrinsic coordinates must enable future state prediction. Specifically, we identify linear dynamics in the matrix : *𝒜* = *A*^nat^ − *A*^act^**K**. This corresponds to minimizing the loss:

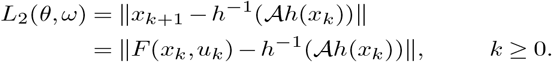

### Linear dynamics

To discover the mapping function *h*, we learn the linear dynamics *A* on the intrinsic coordinates, i.e., *z*_*k*+1_ = *Az*_*k*_. Linear dynamics are achieved via minimizing the following loss:

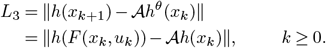

The final loss function is a weighted sum of *L*_1_, *L*_2_, and *L*_3_:

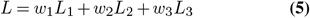

where *w*_1_, *w*_2_, *w*_3_ ∈ ℝ^+^ with *w*_1_ + *w*_2_ + *w*_3_ = 1.

In biological systems, once a wound is created, the natural healing dynamics *A*^nat^ remain relatively stable over time, while the external environment affecting the wound can change frequently. This necessitates a model where *A*^nat^ updates at a slower rate than *A*^act^, and we should consider capturing the temporal relationship of the measurements {*x*_*k*_} and the delayed biological responses to changes in external conditions or treatments. Thus, the *A*^nat^ is updated every *N* samples, and *A*^act^ is updated through all the samples within a window of size *N*, through the attention mechanism detailed in (31).

Note that minimizing the loss *L* in Eq. (5) only gives an estimation of the linear representation for the nonlinear wound healing dynamics, through which an optimal control law can be derived. To derive the real optimal treatment strategy *u*^⋆^ ∈ ϕ^⋆^(*x*), in this paper, we propose a DRL agent to track the reference signal incurred by Eq. (4) whenever it is available and penalize the DRL agent whenever it is not. We show that the control law learned by this DRL agent performs better than the one directly optimizing over a nonlinear system without a mapping in terms of faster convergence and more stability.

### Reinforcement learning algorithm design

In this section, we introduce the use of a DRL algorithm to explore possible treatment policies such that when the wound was treated with this policy, its proceeding state would be as close as to the one generated from linear state with optimal control law. In other word, when the optimal control input Eq. (4) is accessible, the DRL algorithm should be able to exploit its acquired knowledge to generate a policy that closely approximates the resulting nonlinear state to the one achieved through control based on the optimal control. The exploration and exploitation of the DRL algorithm do not require knowledge of either nonlinear or linear dynamics, and thus, it not only alleviates the burden of mathematical interpretation in real-world treatment scenarios but significantly expedites the healing process. To realize this, we first formulate the wound healing dynamics as the Markov decision process (MDP) problem and subsequently solve it using Advantage Actor Critic (A2C) algorithm (32).

Consider a MDP defined by (X, ⋃, *P, r, γ*), where X represents the state space, ⋃ represents the input/action space, *P* represents the transition probability matrix, *r* represents the reward function, and *γ* represents the discount factor. In MDP, an autonomous agent makes sequential discrete-time decisions as time passes. Generally speaking, the MDP problem conforms to the decision-making process of physicians in wound care. Based on the state *x*_*k*_ ∈ X, the agent selects action *u*_*k*_ ∈ ⋃ at time *k*, then it observes the next state *x*_*k*+1_ and receives the reward *r*(*x*_*k*_, *u*_*k*_) ∈ℝ. To collect more state information in wound management, the agent can perform state observation more frequently, such as a state observation every hour and action selection every 20 minutes (11, 29).

The state *x*_*k*_ transits to the next state *x*_*k*+1_ following the transition probability matrix *P* (*x*_*k*+1_| *x*_*k*_, *u*_*k*_), which represents the dynamics of the operating environment. The transition probability matrix satisfies the Markovian (or memoryless) property since a transition to the next state *x*_*k*+1_ depends only on the current state *x*_*k*_ and action *u*_*k*_ rather than a historical series of states and actions. The agent learns the optimal policy ϕ^⋆^ : X *→* ⋃, which maps *x* ∈ X to optimal actions *u* ∈ ⋃ over trial and error interaction with the environment. Nevertheless, the transition probability matrix and the probability distribution of the reward function are generally unknown in reality.

This article designs action, state, and reward based on the characteristics of wound healing. The action *u* ∈ ℝ in the algorithm indicates the the EF (electric field) intensity or the dosage of the drug. The state *x* is an image of the wound. The reward *r* of the algorithm is the exponential of negative Euclidean distance between the next wound image and the image generated by the linear state:

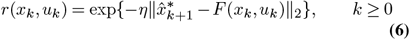

where 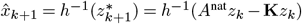, and *η* is a hyperparameter that controls reward’s magnitude.

### Advantage Actor-Critic (A2C) Algorithm Descriptio

This section introduces the A2C algorithm, a reinforcement learning optimization method based on the actor-critic (AC) framework (33). The A2C algorithm differs from value-based algorithms by focusing on policy optimization. It combines two neural networks and employs an advantage function to assess action quality, enhancing the agent’s learning stability. The advantage function reduces action selection variability, leading to more stable learning. Unlike the AC algorithm, the A2C algorithm incorporates an advantage function to stabilize the policy.

The A2C algorithm works by having the actor select actions based on the environment, then, using feedback from the critic, adjust its action selection probability. After the actor completes an action, it returns the new state and value to the critic, which computes the TD (temporal difference) error and updates the action probabilities for the actor. This cycle continues until the termination condition is met.

The key formulas of the A2C algorithm are as follows: We represent the value of action *u* in a given state *x* as *Q*(*x*_*k*_, *u*_*k*_). This paper focuses on the execution of *u* when in state s, and measures the reward of the current action based on the obtained results. *V* (*s*_*t*+1_) represents the value of the subsequent state. The relationship between functions *Q* and *V* is described by the Bellman equation:

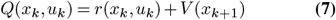

#### Algorithm 1

Closed-loop control of wound healing with A2C.

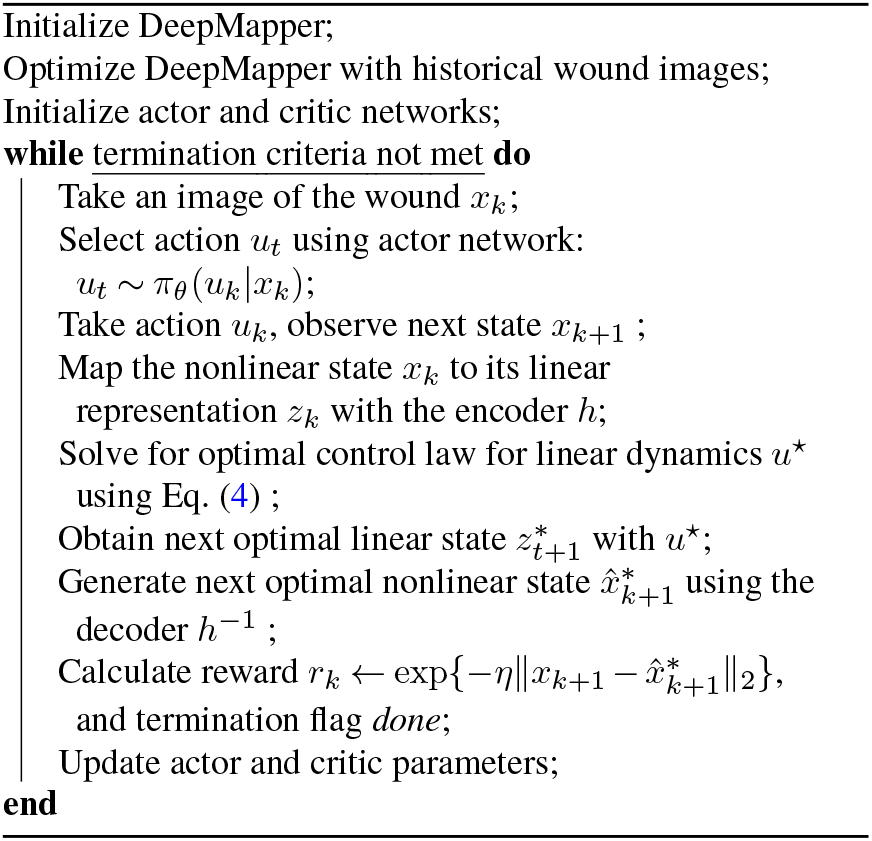

Denote the advantage *A* at state *x*_*k*_ and *u*_*k*_ as:

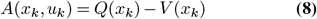

The advantage demonstrates the linear relationship between *Q* and *V*. This equation quantifies the advantages of an action compared with the average value in state *x*, promoting *u* balanced advantage value across the entire strategy. If the advantage value is less than 0, it indicates that this action is inferior to the average and not a good choice. The gradient of the actor-network parameter update is calculate by

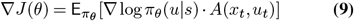

where *J* (*θ*) is the objective function of the actor-network, and *π*_*θ*_ is the probability of action output by the policy network in state 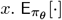 in the formula represents the expected value when selecting an action according to the policy *π*_*θ*_. We update the actor network parameters through Eq. (9). We insert Eq. (7) into Eq. (9) and obtain:

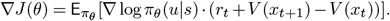

All decision values are calculated in a neural network, making the results more stable.

The steps for obtaining optimal solutions are summarized as follows in Algorithm 1. The optimal of parameters for the neural networks will be obtained by using the Polyak–Ruppert averaging method (34):

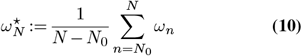

where *N* denotes the total number of updates in the parameters, and the interval [0, *N*_0_] with *N*_0_ *< N* is known as the *burn-in* period; estimates from this period are abandoned to reduce the impact of transients in early stages of the training.

## Experiment Results

Wound-healing studies in humans have been limited by a scarcity of relevant *in vivo* and *in vitro* experimental models and the questionable practice of serial biopsies of human wounds, which are painful and may cause infection or scarring. Mouse models have an advantage because they are small, easy to handle, and relatively cheap. However, a high degree of phenotypic variability in skin exists between animal species, and the impact of these differences on wound healing has been recognized since at least 1915 (35). Therefore, an accurate model of human wound healing should use an animal with similar skin characteristics.

Porcine models have emerged as promising models to study wound healing, with over 1500 publications on the psychophysiology of various wound types. An advantage of using pigs is that they are anatomically and physiologically similar to humans, and have been used to study many other diseases (36), such as diabetes, cardiovascular diseases, and infections. Similarity between pig and human skin also make pigs an appropriate model for cutaneous wound healing. Like humans, they have a relatively thick epidermis, distinct rete pegs, dermal papillae, and dense elastic fibers in the dermis (37, 38).

In this paper, we conducted two numerical studies: one on using mouse data, and one on *in vivo* porcine experiments.

### Mouse Data Simulation

The mouse image dataset used in this work was created as described in (39, 40), from which we generated circular wound-only crops. The dataset includes wounds from eight mice (four from one cohort and four from another distinct cohort) imaged daily over sixteen days. Each mouse had two wounds, one on the left side and one on the right side. This process resulted in a total dataset of 256 images (8 mice × 2 wounds × 16 days). We split the dataset into training (224 images) and testing (32 images) subsets.

An example trajectory of testing images is shown in Fig. 2 (a). The natural healing process of mouse wounds typically spans approximately 10 to 11 days. In our study, we employed *A*^nat^ and *A*^act^ as detected by the DeepMapper model. The objective was to demonstrate whether the proposed algorithm can accelerate the wound-healing process in mice.

**Fig. 2.**
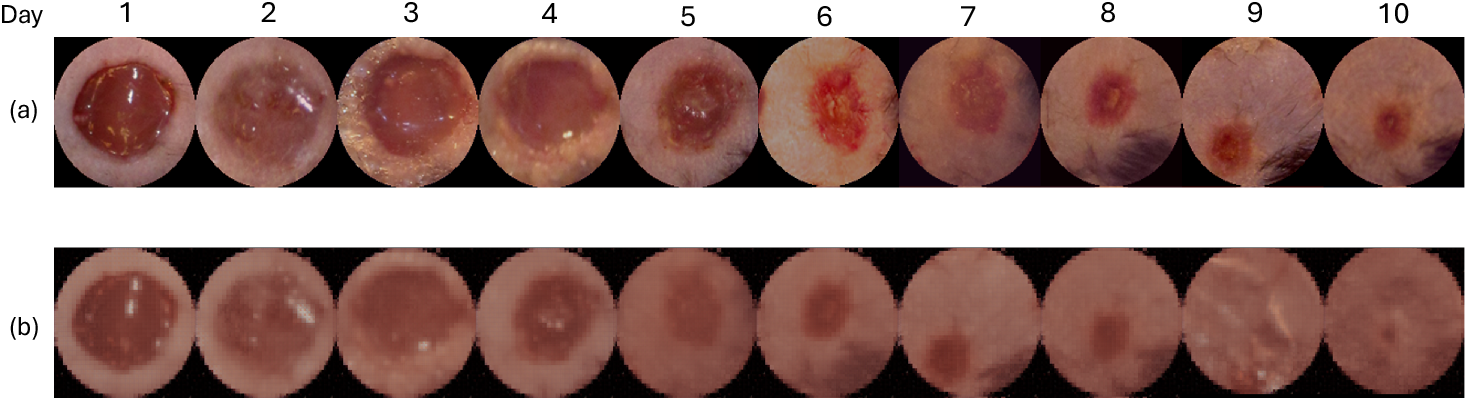
Trajectories of wound images. (a). One trajectory of mouse wound images in the testing dataset. It would take around 10 to 11 days for the mouse wound to heal. (b). One trajectory of generated mouse wound images using control policy given by the DRL agents. On day 9, there’s no obvious wound regions in the generated images.

Since no treatment was applied to the mouse wounds during data collection, *A*^nat^ will remain unchanged. However, the components 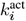 and 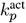 in *A*^act^ were modulated to new values, denoted as 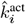 and 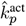, respectively. This modulation was achieved via a predefined function that simulates the effects of electrical fields and fluoxetine, described by the following equations:

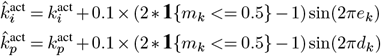

The goal was to optimize the control input *a*_*k*_ := [*e*_*k*_, *d*_*k*_] such that the cumulative time with wound metrics *m*_*k*_ *≤* 1 is minimized.

To achieve this, we trained DRL agents to derive the control policy using the reward function defined in Eq. (6) and *A*^nat^ and *A*^act^ predicted by the DeepMapper from training images. The DRL agents were tested every 50 training episodes on the testing images, with 100 independent runs conducted using 100 different random seeds.

The results are presented in Fig. 3, showing that the control policy derived by the DRL agents reduced the wound healing time to approximately 8.5 days, compared to 10.33 days for normal healing (17.71% faster). Moreover, the generated wound images in Fig. 2 (b) show no visible wound regions by day 9, further validating the efficacy of the proposed method. Additionally, we compared this control policy with a policy optimized using a simpler reward function that directly targeted minimizing wound healing time without employing the leader-follower tracking strategy proposed in this paper:

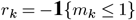

**Fig. 3.**
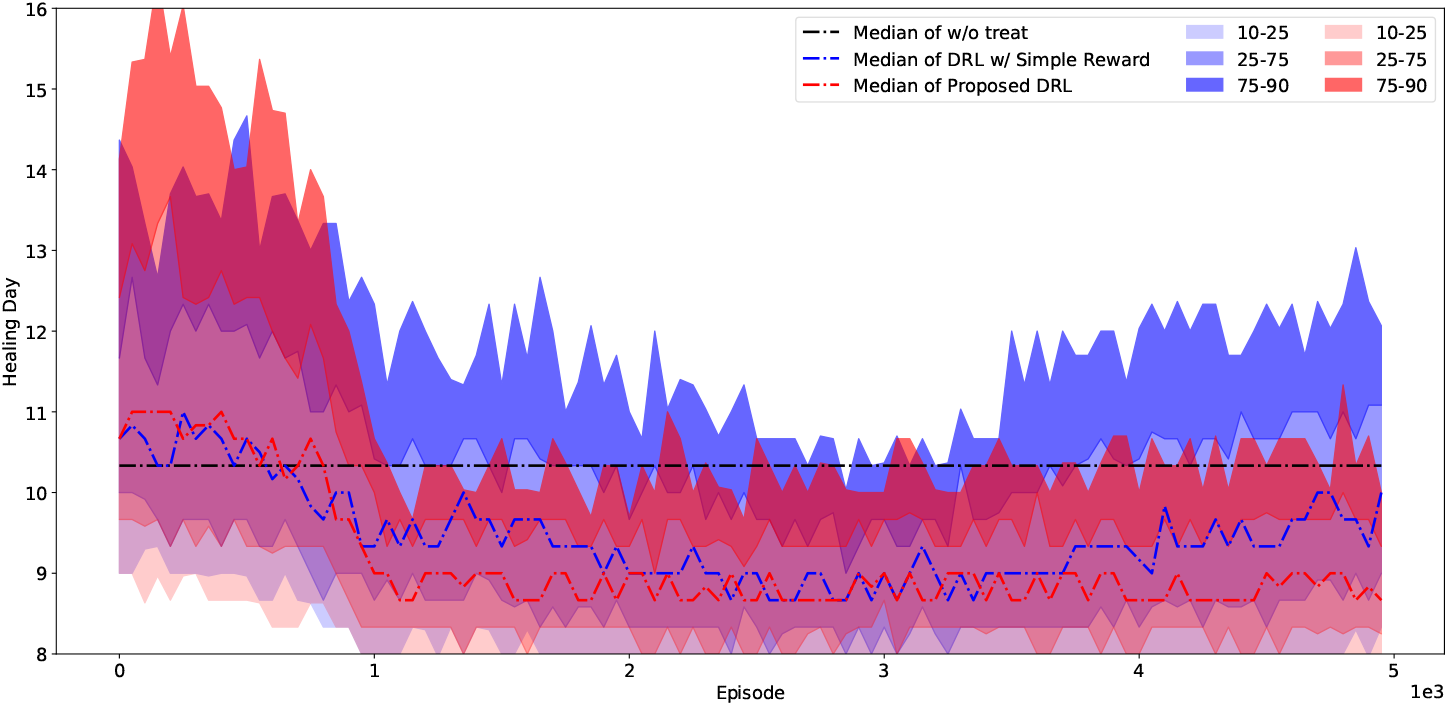
Learning curves of DRL agents over mouse dataset with a predefined function of treatment effect, shown by percentile. Without any treatments, it would take 10.33 days for one mouse wound to heal. Our proposed method, shown in red curves, results a treatment policy that is more stable than the DRL agents without the leader-follower strategy.

The blue curves in Fig. 3 illustrate that this alternative approach not only converged more slowly than the proposed method but also exhibited higher variance. These findings highlight the effectiveness and robustness of the proposed DRL-based control strategy in expediting the wound healing process.

### In vivo Application

This work was further integrated into a 22-day *in vivo* porcine experiment in (22). A total of 6 to 10 wounds, each 20 mm in diameter, were created on the dorsum of a pig. These wounds were interfaced with a bioelectronic device with 8 treatment delivery channels: 4 channels were designed for fluoxetine delivery, and the rest for application of an electric field. Both the fluoxetine dosage and electric field intensity were controlled via electric currents in the corresponding channels. The dosage of fluoxetine delivered from each channel was controlled by the current applied. The wounds were also interfaced with a camera that takes real-time images of the wound, based on which the proposed algorithm predicts wound stages and the DRL agent makes decisions on the applied currents to control the dosage or intensity.

The real-time imaging system in (41, 42) captured wound images every 2 hours throughout the experiment. We fine-tuned DeepMapper to work with the device images. An image of the device and a sample wound image captured by the device are shown in Fig. 5. Note that they are substantially different than standard *in-vivo* wound images, which have wound edge information and different color spectrum. To pre-process the images, we subtracted [108.16, 61.49, 55.44] from the pixel values of the RGB channels in wound image to reduce redness, and we down-sample the image to shape (H: 128, W: 128, 3). The down-sampling method we are using is called Lanczos algorithm (43). Finally, we normalize pixel values to have zero mean and range from 0 to 1; The processed wound images were passed into the DeepMapper to predict the wound’s healing stage as well as the linear dynamics, based on which optimal reference signals were calculated via Eq. (4) and the DeepMapper’s decoder.

Fig. 4a shows the wound stage prediction and solutions to predicted linear dynamics. In the ideal case, predicted wound stage and the solution to the linear dynamics should align with each other. However, since DeepMapper was pretrained on historical wound images, its prior knowledge about wound evolution tends to dominate early predictions, leading to deviations from the actual linear evolution. For instance, we observed a mean squared error (MSE) of approximately 69.15% on the first day predictions. As more data is collected over time, the dotted curves in Fig. 4a increasingly align with the linear dynamics solutions, with the MSE reducing to 3.71% by the end of the first day.

**Fig. 4.**
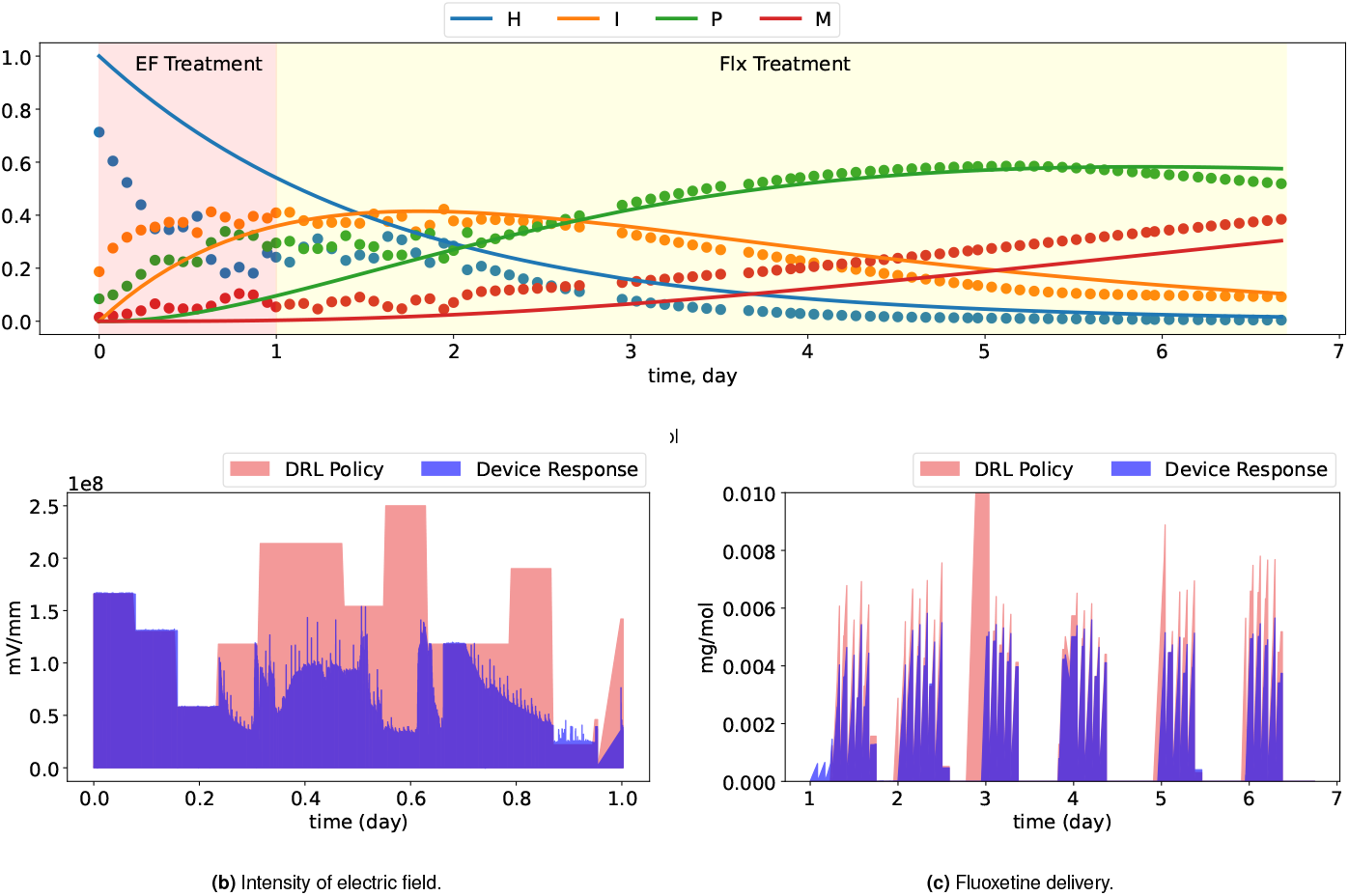
Plots of first 7-day wound stage predictions, solution to learned linear dynamics, and treatment policies given by the DRL agents across the 22-day porcine experiment.

**Fig. 5.**
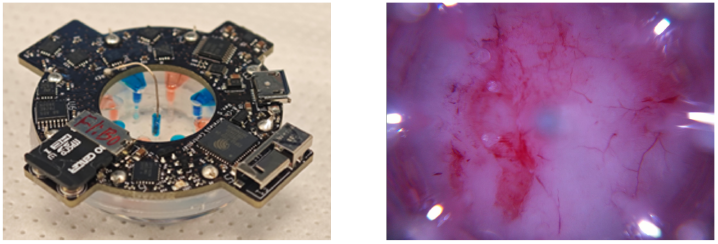
An image of EF+Flx combo actuators used in the *in vivo* experements to deliver EF and fluoxetine (shown on the left), and a sample wound image captured by the device camera (shown on the right).

Two DRL agents then used would images to make dynamic adjustments to the currents, optimizing either fluoxetine dosage and electric field intensity so that the cumulative reward Eq. (6) is maximized. In our experiment, only one DRL agent was active at a time, meaning that when DRL agent for electric field was active, all other channels associated with fluoxitine delivery were shut off. Initially, we used the agent controlling the electric field; once the predicted healing probability reached approximately 40% and started to decrease, we switched to the DRL agent for fluoxetine delivery.

On day 0, wounds were created. These wounds were categorized into two groups: one actuator-camera combination DRL treated wounds, and one camera-only control wound. Devices were applied to the wounds while the pig was under anesthesia. The devices operated in closed-loop controlled delivery mode for 22 hours each day. Power banks were replaced daily during feeding. On day 3, the devices were replaced with new ones. The DRL agent for application of electric field was initiated on day 0, and switched to DRL agent for fluoxetine in the middle of day 1, continuing fluoxetine treatment until day 7. The treatment strategies for intensity of electric field and dosage of fluoxetine are illustrated in Fig. 4b and Fig. 4c.

On day 7, devices were removed and replaced with standard-of-care treatments, and the experiment continued until harvest on day 22. After that, the animal was sacrificed, and tissue samples were collected for analysis.

Fig. 6 shows wound progression for the DRL treated wound and control wound. Our primary goal is to double the healing rate compared to the natural wound healing process. The DRL treated wound had achieved 42.26% progress toward full closure—nearly 40% faster than the control wound, which had reached only 30.12% progress, based on the wound stage prediction by DeepMapper. This result suggests that the proposed algorithm provided a treatment strategy that effectively accelerated the healing process.

**Fig. 6.**
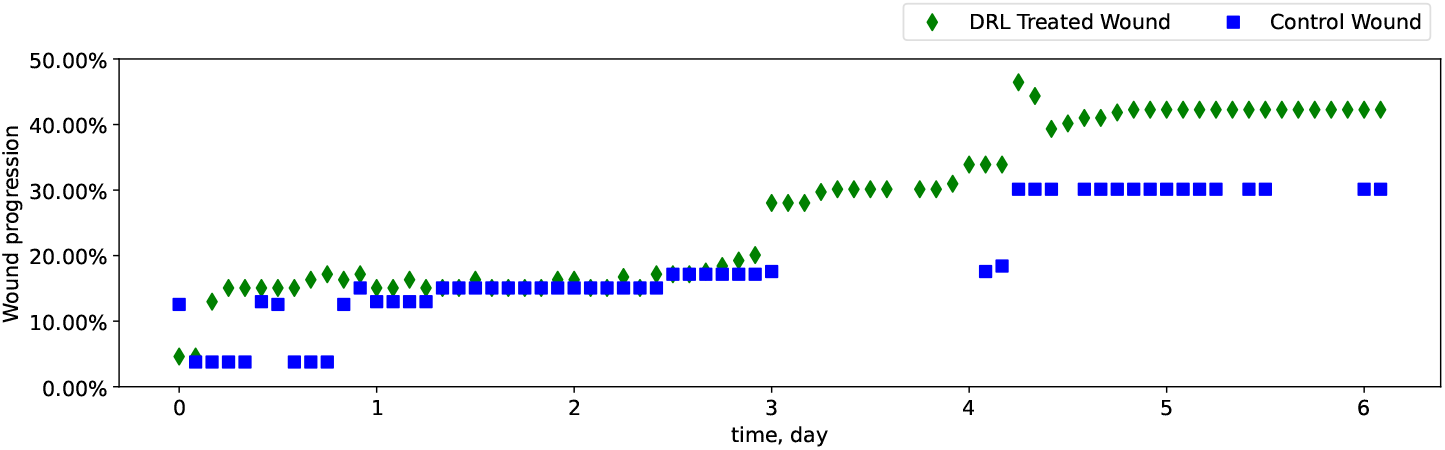
Wound progression tracking plot based stage predictions of the DeepMapper. The DRL treated wound shows 40.26% progress toward full closure - nearly 40% progress faster than the control wound, which had reached only 30.12% progress.

## Discussion

The advent of new biotechnologies has introduced an array of sensors for biological systems, addressing a critical need in medicine to integrate these sensors into closed-loop control systems. However, the inherent complexity of biological processes poses a challenge for formulating precise mathematical models. Consequently, there is a growing demand for control algorithms that can operate effectively without relying on exact models. Although sensors provide valuable insights, their measurements only offer a partial view of the underlying dynamics of real biological systems.

Wound healing exemplifies a nonlinear biological process, where different cell types play distinct roles at various stages. In our study, we assume the availability of a sensor capable of reflecting wound stages, as suggested in (29), with the sensor data being approximately modeled by a linear system of ordinary differential equations (ODEs).

Despite the convenience of using linear models, discrepancies between sensor measurements and the actual biological processes involved in wound healing reveal the limitations of linear systems in capturing the underlying nonlinear dynamics.

Nevertheless, we demonstrate that nonlinear systems can be effectively monitored and controlled using observations derived from linear approximations. This approach shows promise for a wide range of nonlinear biological processes where sensors exist, but accurate mathematical models are difficult to construct.

It is also important to note that more advanced deep reinforcement learning (DRL) algorithms can further enhance the control strategies and improve sample efficiency. For instance, in large state and action spaces, Actor-Critic (A2C) methods may suffer from slow convergence and susceptibility to local minima. More advanced approaches, such as Asynchronous Advantage Actor-Critic (A3C) (32), can leverage parallelism to mitigate these issues. However, these algorithms still struggle with convergence due to the non-convex nature of their objective functions. The integration of convex optimization techniques, as explored in (44), holds potential for overcoming this limitation.

The *in vivo* experimental results presented in this paper, specifically the acceleration of wound healing, were measured using predictions from DeepMapper. While our findings suggest significant differences between treated and control wounds, further studies are required. Additional experiments and data collection are necessary to fine-tune the deep learning models used in the proposed approach. Moreover, biological analysis of the wounds by experts is needed to substantiate the results. However, due to the prolonged nature of the wound-healing process, these experiments can be both time-consuming and expensive.

Our primary objective is to propose an adaptive learning framework that combines deep learning, optimal control, and deep reinforcement learning to accelerate wound healing. Future work will focus on designing more efficient DRL algorithms tailored to these goals.

## Conclusion

In this paper, we propose an adaptive closed-loop control framework for a nonlinear dynamical system. The controller integrates deep learning, optimal control, and reinforcement learning, aiming to accelerate nonlinear biological processes such as wound healing without the need for mathematical modeling. We have demonstrated that the proposed method not only significantly improves wound healing time but also addresses safety concerns and reduces drug usage.

In the future, we aim to enhance the algorithm’s performance by reducing the action space and minimizing the selection of prohibited actions, which requires the analysis of biological effects of each action been executed, thereby improve its interpretability. Additionally, we will explore the application of alternative reinforcement learning algorithms to achieve significant enhancements (32, 44).

## ACKNOWLEDGEMENTS

This study was supported by the SciAI Center and funded by the Office of Naval Research (ONR) under Grant Number N00014-23-1-2729 and the DARPA Biotechnologies Office (DARPA/BTO) under Cooperative Agreement Number DC20AC00003. The porcine wound images were captured by Da Vinci Services.

